# Multimodal Imaging Investigation of Rich Club Alterations in Alzheimer’s Disease and Mild Cognitive Impairment: Amyloid Deposition, Structural Atrophy, and Functional Activation Differences

**DOI:** 10.1101/2023.08.21.554083

**Authors:** Sebastian Markett, Ole J. Boeken, Olga A. Wudarczyk, the Alzheimer’s Disease Neuroimaging Initiative

**Affiliations:** Humboldt-Universität zu Berlin, Berlin, Germany; Charité-Universitätsmedizin, Department of Neurology and Experimental Neurology, Berlin, Germany; International University of Applied Sciences, Berlin, Germany

**Author notes:** corresponding author’s address: Dr. Sebastian Markett, Humboldt Universität zu Berlin, Unter den Linden 6, 10099 Berlin, Germany. Data used in preparation of this article were obtained from the Alzheimer’s Disease Neuroimaging Initiative (ADNI) database (adni.loni.usc.edu). As such, the investigators within the ADNI contributed to the design and implementation of ADNI and/or provided data but did not participate in analysis or writing of this report. A complete listing of ADNI investigators can be found at: http://adni.loni.usc.edu/wp-content/uploads/how_to_apply/ADNI_Acknowledgement_List.pdf.

## Abstract

Alzheimer’s disease (AD) is characterized by significant cerebral dysfunction, including increased amyloid deposition, gray matter atrophy, and changes in brain function. The involvement of highly connected network hubs, known as the “rich club,” in the pathology of the disease remains inconclusive despite previous research efforts. In this study, we aimed to systematically assess the link between the rich club and AD using a multimodal neuroimaging approach.

We employed network analyses of diffusion MRI, longitudinal assessments of gray matter atrophy, amyloid deposition measurements using PET imaging, and meta-analytic data on functional activation differences. Our study focused on evaluating the role of both the core and extended rich club regions in individuals with Mild Cognitive Impairment (MCI) and those diagnosed with Alzheimer’s Disease.

Our findings revealed that rich club regions exhibited accelerated gray matter atrophy and increased amyloid deposition in both MCI and Alzheimer’s Disease. Importantly, these regions remained unaffected by altered functional activation patterns observed outside the core rich club regions. These results shed light on the connection between two major AD biomarkers and the rich club, providing valuable insights into AD as a potential disconnection syndrome.

## 1. Introduction

Alzheimer’s disease is a neurodegenerative disorder characterized by a progressive decline in cognitive function, leading to significant emotional, social, and societal burdens for patients, families, and caregivers (Reitz et al., 2011; Weller & Budson, 2018). At the neurobiological level, the disease is marked by cerebral atrophy, including volume loss, morphological changes, and cortical thinning (Pini et al., 2016). Another key feature is the accumulation of amyloid-beta plaques, abnormal protein fragments that disrupt cell function and communication between neurons (Blennow et al., 2006; Hardy & Selkoe, 2002; Liu et al., 2013; Mawuenyega et al., 2010). Additionally, the presence of tau pathology, characterized by abnormal modifications of the tau protein and the formation of neurofibrillary tangles within neurons, further impairs neuronal function (Ittner & Götz, 2011; G. Lee & Leugers, 2012).

An influential hypothesis suggests that Alzheimer’s disease can be seen as a disconnection syndrome, affecting the brain’s network infrastructure (Brier et al., 2014; Delbeuck et al., 2003; Geschwind, 1965; Wang et al., 2015). This disruption leads to disorganized network configurations, with some brain regions being more vulnerable to pathological changes than others (Brier et al., 2014; Fathian et al., 2022; Wang et al., 2015). These particularly vulnerable regions often correspond to highly connected network hubs (Crossley et al., 2014), which play crucial roles in information processing (van den Heuvel & Sporns, 2013). Within the cerebral cortex, hubs are found in heteromodal areas of the association cortices and are susceptible to amyloid-beta deposition, atrophy, and disruption of activity and metabolism in Alzheimer’s (Buckner et al., 2009; Dai et al., 2015; Stam et al., 2009; Yu et al., 2017).

The human brain’s densely connected hub regions are thought to form a centrally connected ‘rich club’ (van den Heuvel & Sporns, 2011). This rich club exhibits high interconnectivity among its hub regions, serving as a central backbone for global communication and information integration (van den Heuvel et al., 2012). However, due to its dense connectivity and high metabolic demands, the rich club is also susceptible to pathology (Griffa & Van den Heuvel, 2018). While reduced integrity of white matter connections between rich club regions has been observed in several brain disorders (de Lange et al., 2019), its involvement in Alzheimer’s disease is still inconclusive.

Studies investigating rich club organization in Alzheimer’s disease have yielded inconsistent findings, thereby warranting further investigation. Some studies have reported preserved rich club organization in individuals with Alzheimer’s disease (Daianu et al., 2015; Ma et al., 2022), while others have observed alterations in the organization, either an increase (Daianu et al., 2016; W. J. Lee et al., 2018; Yan et al., 2018) or decrease (Drenthen et al., 2022; Fathian et al., 2022; Xue et al., 2020). These discrepancies in findings may be attributed to methodological variations across studies such as different analytic approaches and imaging techniques (Wu et al., 2019). While the literature presents divergent findings on the connectivity between rich club regions in Alzheimer’s disease, consensus emerges when examining feeder connections, which link peripheral (i.e. non rich club member) brain regions to the rich club. Multiple studies consistently report a decrease in connectivity of feeder connections in the context of Alzheimer’s disease, indicating their potential involvement in the disease’s pathophysiology (Cao et al., 2020; Daianu et al., 2016; W. J. Lee et al., 2018; Mirza-Davies et al., 2022).

Despite evidence suggesting a spatial co-localization of Alzheimer-related pathology with network hubs in the human brain, the specific role of the rich club in Alzheimer’s disease remains poorly understood. To address this knowledge gap, we aim to investigate the potential involvement of rich club regions in Alzheimer’s disease by examining the progression of atrophy, amyloid deposition, and functional disturbances. To accomplish this, we adopt a multimodal neuroimaging approach, integrating structural connectivity data from a cohort of healthy elderly participants with case-control data of patients with Alzheimer’s disease and mild cognitive impairment (MCI) that include longitudinal assessments of gray matter atrophy in magnetic resonance imaging (MRI) data, assessments of amyloid deposition in positron emission tomography (PET) data, and meta-analytic indicators of functional disturbances. To obtain a comprehensive understanding of the rich club in Mild Cognitive Impairment and Alzheimer’s Disease, we extend our investigation beyond the narrow-sense core rich club and include an analysis of the larger extended rich club, considering that the rich club concept encompasses both a specific subset of highly interconnected hub regions and a broader principle of network organization, where also less heavily-connected regions exhibit preferential connections beyond their individual connectivity degree. For both the core rich club and for the extended rich club, we ask whether rich club regions show accelerated gray matter atrophy, increased amyloid deposition, and more signs of functional disturbances than the remaining peripheral regions in patients relative to healthy controls.

## 2. Methods

Part of the data used in the preparation of this article were obtained from the Alzheimer’s Disease Neuroimaging Initiative (ADNI) database (adni.loni.usc.edu). The ADNI was launched in 2003 as a public-private partnership, led by Principal Investigator Michael W. Weiner, MD.

The primary goal of ADNI has been to test whether serial magnetic resonance imaging (MRI), positron emission tomography (PET), other biological markers, and clinical and neuropsychological assessment can be combined to measure the progression of mild cognitive impairment (MCI) and early Alzheimer’s disease (AD). For up-to-date information, see www.adni-info.org.

The present investigation combines neuroimaging data from three different cohorts: a) Structural MRI for the assessment of gray matter atrophy in patients with AD, MCI, and healthy controls from the ADN1 cohort (Petersen et al., 2010), b) florpetapir PET data for the assessment of amyloid depositions in patients with AD, MCI, and healthy controls from the ADNI-GO/2 cohort, and c) structural connectivity data from the 10kin1day cohort (van den Heuvel et al., 2019) for building a reference connectome from elderly healthy controls. These data are complemented by meta-analytic reverse inference decoding in *Neurosynth* (Yarkoni et al., 2011) to address functional disturbances in AD and MCI. All three data sets and the meta-analytic approach will be described in the following. We combined data from all imaging modalities at the level of 82 cortical and subcortical regions of interest (ROIs) as described in the Desikan-Killiany atlas (Desikan et al., 2006).

### 2.1 Structural MRI data

We included N = 424 participants (n = 224 males, n = 200 females) from the ADNI data set with at least two available MRI scans (at screening and at twelve months follow-up) that had been processed with Freesurfer’s longitudinal pipeline. The sample included n = 72 patients with Alzheimer’s disease (age M = 73.93, SD = 7.99, n = 34 male, n = 38 female), n = 215 patients with MCI (age M = 74.04, SD = 7.24, n = 124 male, n = 91 female), and n = 137 healthy control participants (M = 74.86, SD = 4.85, n = 66 male, n = 71 female). All structural MRI data were acquired at 1.5T.

### 2.2 PET data

We included N = 363 participants from the ADNI-GO/2 cohort who had undergone structural MRI imaging at 3T at baseline and at least one session of PET imaging with florbetapir (^18^F-AV-45) as radionucleotide. In case that more than one PET session was available, we selected the session closest to the baseline MRI scan. The sample included n = 102 patients with Alzheimer’s disease (age M = 74.23, SD = 8.14, n = 57 male, n = 45 female), n = 119 patients with MCI (age M = 72.54, SD = 7.92, n = 61 male, n = 58 female), and n = 142 healthy controls (age M = 73.43, SD = 6.22, male n = 72, female n = 70).

### 2.3 Connectome data

We used connectome data from the openly available 10kin1day data (van den Heuvel et al., 2019). This data set contains structural connectome data from 8,168 participants. From this dataset, we selected connectome matrices from healthy (i.e. non patient) participants aged 55 to 90 years to match the age range from the ADNI data sets. This resulted in a final sample of N = 865 participants (n = 417 males, n = 446 females, n = 2 no gender specified) from eight age groups (50-55 n = 224, 55-60 n = 163, 60-65 n = 121, 65-70 n =186, 70-75 n = 95, 75-80 n = 51, 80-85 n = 21, 85-90 n = 4). Details on the processing pipeline and quality control are given in van den Heuvel et al. (2019). In brief, connectomes were assembled by first obtaining a cortical and subcortical gray matter parcellation from running T1-weighted structural images through Freesurfer (Fischl et al., 2004) and then collating the resulting parcellation with DTI (diffusion tensor imaging) data. Diffusion data were first corrected for susceptibility and eddy current distortions. Then each voxel’s main diffusion direction was obtained via robust tensor fitting. Large white matter pathways were formed by deterministic fiber tractography (Mori et al., 1999). Fiber streamlines were propagated along each voxel’s main diffusion direction after originating from eight seeds evenly distributed across each white matter voxel until a stopping criterion was met (hitting a voxel with FA <.1, a voxel outside the brain mask, or making a turn of > 45 degrees). A pair of regions from the gray matter parcellations was considered connected when both regions were touched by a reconstructed streamline. Connections were weighted with the total number of streamlines that touched both ROIs. This resulted in one weighted and undirected connectome matrix for each individual.

### 2.4 Network and rich club analysis

Network analysis and extraction of network parameters were performed in Matlab. We created a weighted 82*82 group connectome adjacency matrix that contained connections present in at least 60% of all participants, weighted by the mean number of streamlines computed per connection across all participants (de Reus & van den Heuvel, 2013). We followed standard procedures for rich club analysis (Riedel et al., 2022; van den Heuvel et al., 2013) based on code from the Brain Connectivity Toolbox (BCT, Rubinov & Sporns, 2010). A network is said to have rich club properties when high degree nodes show a higher level of interconnectedness than expected from their high degree alone (van den Heuvel & Sporns, 2011), across a range of degree thresholds. The rich club regime was established as follows: We first computed the weighted rich club coefficient (using the BCT function rich_club_wu.m) across the full range of levels k from the network’s degree distribution (k=1,..,35). Because high degree nodes have a high likelihood to connect to other high degree nodes by chance alone, it is necessary to establish that the empirical level of interconnectedness exceeds the level of interconnectedness in random network null models. We created 10,000 null networks by reshuffling all connections in the matrix, effectively destroying network topology. We used the BCT function randmio_und.m that preserves the degree distribution of the network. Each connection was rewired 10 times. Standardized rich club coefficients were then obtained by dividing empirical coefficients by coefficients from all 10,000 iterations of the null model across the full range of k. We determined the rich club regime as the largest series of subsequent k, where the empirical rich club coefficient was larger than the rich club coefficient in 95% of all null networks. We assigned nodes to the rich club when their nodal degree was equal to or larger than the k-value where the normalized rich club coefficient was maximal. We complemented this definition by a second analysis of an ‘extended rich club’ which included all brain regions with nodal degree equal to or larger than the k-value that marked the starting point of the rich club regime.

### 2.5 Atrophy analysis

Atrophy for each participant and for each of the 82 regions of interest was computed by subtracting mean gray matter volume at follow up from mean gray matter volume at baseline (12 months earlier). We computed mean atrophy for rich club regions and for peripheral (i.e. non-rich club) regions.

### 2.6 PET analysis

Regions of interest were delineated at baseline by running structural images through Freesurfer. Mean florbetapir update was computed for each of the 82 regions of interest. Mean uptake was standardized with the size of the respective ROI (mean gray matter volume of ROIs from the baseline MRI scan). For each level K, standardized regional uptake ratio value (SUV) of florbetapir PET was calculated across rich club and non-rich club regions by dividing the sum of the florbetapir-volume-products across regions by the sum of the volumes across regions.

### 2.7 Meta-Analysis

Functional characterization of all 82 regions of interests was achieved via meta-analytic reverse inference decoding, using the NeurosynthDecoder as implemented in the Neuroimaging Meta-Analysis Research Environment (NiMARE; Salo et al., 2022), following our standard protocols (Boeken et al., 2022; Boeken & Markett, 2023). Reverse inference decoding seeks to establish associations between a given term (such as ‘alzheimer’ or ‘mild cognitive’) and regional activity by determining the relative over-representation of these term in studies reporting activation at a particular location. To this end, the term-specific activations are compared to the entire Neurosynth database which contains functional activation coordinates from 14,371functional neuroimaging studies, annotated with around 3000 terms, covering various categories such as psychological constructs (e.g., ‘memory’), disease related-terms (e.g.,’alzheimer’, ‘mild cognitive’), and anatomical terms (Yarkoni et al., 2011). We queried the database with all 82 regions of interest and searched the resulting datasets for the terms “alzheimer”, “alzheimer disease”, “mild cognitive”, “mci” (i.e., both as for mild cognitive impairment) and “cognitive impairment”. We then calculated a two-way chi-square statistic for each region to assess the statistical independence of the label’s presence and the term selection, using the Benjamini-Hochberg procedure to control the false discovery rate. For all significant region-term associations, we then calculated the Bayes Factor as the ratio between the posterior odds and the prior odds (Goodman, 1999; Poldrack, 2006), to obtain an interpretable measure of association strength of the term given a possible activation within the given region of interest.

### 2.8 Statistical Analysis

Atrophy and amyloid data were analyzed by analyses of variance, treating rich club as within-subject factor (rich club vs. periphery), group as between-subject factor (Alzheimer, MCI, healthy controls), and participants’ age as covariate. Significant interactions between rich club and group were further explored by post-hoc comparisons that focused on two groups only (healthy controls vs. MCI, healthy controls vs. Alzheimer, MCI vs. Alzheimer).

For each meta-analytis, we calculated Bayes factors for rich club and non-rich club regions by averaging the regional Bayes factors for both sets of brain regions. Statistical significance of the association between the rich club and functional activation differences was examined through a series of χ^2^-tests, examining whether brain regions with meta-analytical activation differences were more likely to be rich club members or not.

### 2.9 Data and Code Availability

Access to atrophy and amyloid data can be obtained though ADNI. Access to the 10k1day connectome data can be obtained through dutchconnectomelab.nl. Raw data from the meta-analysis, as well as all analysis code to perform the connectome analyses, the meta-analyses, and to re-create the figures can be obtained from our Github repository (https://github.com/markett-lab/AlzheimerRichClub).

## 3. Results

### 3.1 Rich Club

We confirmed rich club organization in the weighted reference connectome (figure 1A), starting from nodes with degree >=12 up to degree >=27 (p <.05, FDR-corrected, figure 1B). The peak of the normalized rich club curve at k>=26 revealed a rich club of seven nodes (8.54%) for which the rich club effect was maximal (figure 1C, red nodes). These regions included: the putamen bilaterally, superior frontal and superior parietal cortex bilaterally, and the left insula. The rich club regime, however, started already at k>=12, identifying 46 additional regions in the extended rich club (53 nodes in total, 64.64%, figure 1C, all highlighted nodes in red and blue).

**Figure 1.**
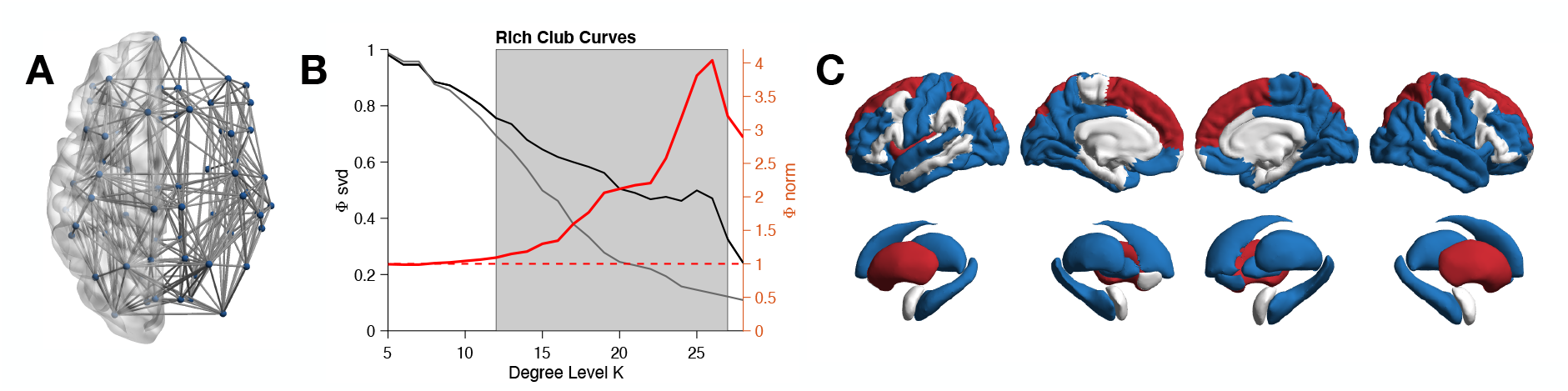
(A) Structural reference connectome based on the standard Freesurfer atlas, (B) Rich club curves (B) for the empirical (black), null model (gray), and normalized (red) rich club coefficients in the reference connectome. The shaded area indicates the rich club regime where the normalized rich club coefficient was significantly larger than then 1 (dashed line). (C) Rich club regions: the red regions are the rich club according to the maximal rich club coefficient (k>=26), the blue regions are the additional members of the extended rich club (k>=12).

### 3.2 Atrophy

We assessed gray matter atrophy of the rich club and the extended rich club longitudinally in patients with Alzheimer, MCI, and healthy controls (Figure 2). Disease status was related to gray matter atrophy, irrespective of participants’ age (main effect: F(2,420)=7.8, p<.001, η^2^=.036): Alzheimer patients showed higher atrophy than MCI patients (F(1,284)=6.763, p=.01, η^2^=.023), and MCI patients showed higher atrophy than healthy individuals (F(1,349)=4.183, p=.042, η^2^=.012). Disease status was differentially related to atrophy of rich club regions (interaction: F(2,420)=3.170, p=.043, η^2^=.015). Relative to healthy controls, Alzheimer patients showed more pronounced atrophy in their rich club (F(1,206)=6.387, p=.012, η^2^=.03). There was no such effect in MCI patients (F(1,349)=1.466, p=.227, η^2^=.004). For the extended rich club, disease status was also differentially related to atrophy (interaction: F(2,420)=7.99, p<.001, η^2^=.037). Relative to healthy controls, both Alzheimer patients (F(1,206)=16.414, p<.001, η^2^=.074) and MCI patients showed higher atrophy in extended rich club regions (F(1,349)=5.039, p=.025, η^2^=.014).

**Figure 2.**
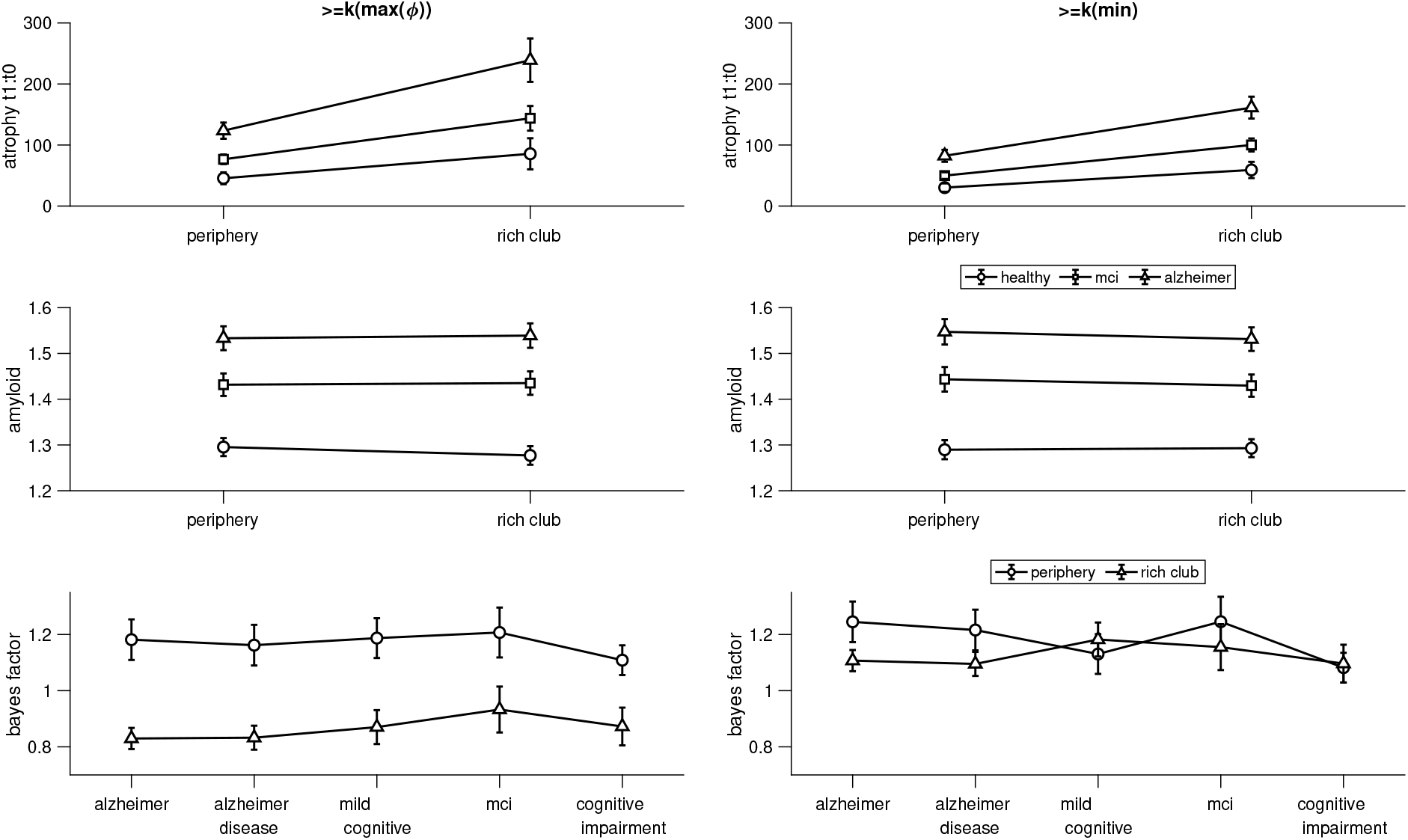
Atrophy (top row), Amyloid (middle row), and functional (bottom row) results for rich club and peripheral regions in Alzheimer patients, patients with MCI, and healthy controls. The left panels use a rich club definition based on the maximal rich club coefficient. The right panels use an extended rich club definition, according to the rich club regime.

### 3.3 Amyloid

We assessed amyloid load of the rich club and the extended rich club in patients with Alzheimer, MCI, and healthy controls (Figure 2). Disease status was related to amyloid load, irrespective of participants’ age (main effect: F(2,359)=28.885, p<.001, η^2^=.139): Alzheimer patients showed higher amyloid load than MCI patients (F(1,218)=8.457, p=.004, η^2^=.037), and MCI patients showed higher amyloid load than healthy individuals (F(1,258)=22.698, p<.001, η^2^=.081). In the entire sample, we observed slightly elevated amyloid load in peripheral regions relative to rich club regions (main effect: F(1,359)=4.215, p=.041, η^2^=.012). This effect, however, was qualified by a significant interaction with disease status F(2,359)=11.983, p<.001, η^2^=.063). The higher amyloid load in peripheral regions was only seen in controls but not in MCI patients (interaction: F(1,258)=16.625, p<.001, η^2^=.061) and not in Alzheimer patients (interaction: F(1,241)=19.971, p<.001, η^2^=.077). Amyloid load differed between the extended rich club and the periphery depending on disease status (interaction: F(2,359)=11.9, p<.001, η^2^=.062). Relative to healthy controls where amyloid load did not differ between extended rich club and peripheral regions, MCI patients (interaction: F(1,258)=16.7, p<.001, η^2^=.061) and Alzheimer patients (interaction: F(1,241)=21.582, p<.001, η^2^=.082) had a higher amyloid load in the periphery.

### 3.4 Meta-Analysis

We queried the Neurosynth database for meta-analyses of terms related to Alzheimer and MCI, collating results from case-control comparisons across the functional imaging literature (Figure 2). Bayes factors – averaged across all rich club and all peripheral regions – suggest that peripheral regions are slightly more likely to be activated than core rich club regions when comparing patients to healthy controls. No such difference was seen for the extended rich club. Across all brain regions, there was no significant difference in meta-analytically associated brain regions between rich club regions and the periphery (χ^2^-tests, all p>.05).

## 4. Discussion

We investigated the potential role of the rich club - a group of highly interconnected and central hub regions in the connectome - in Alzheimer’s disease (AD) and mild cognitive impairment (MCI). To achieve this, we combined structural connectivity data with longitudinal assessments of gray matter atrophy, amyloid deposition measurements using PET imaging, and meta-analytic data on functional activation differences.

Next to the core rich club, which included the putamen, superior frontal and parietal cortices, and the insula, in accordance with the previous literature (Collin, Sporns, et al., 2014; Markett et al., 2017; van den Heuvel et al., 2013; van den Heuvel & Sporns, 2011; Verstraete et al., 2014), we also examined an ‘extended rich club’, following previous observations that rich club organization is a general principle of brain networks and not of a few high-degree nodes only (Griffa & Van den Heuvel, 2018; Riedel et al., 2022; van den Heuvel & Sporns, 2011). The extended rich club included all brain regions that demonstrated stronger interconnectedness than expected based on the number of their connections alone, and thus fell under the network’s rich club regime. We found that approximately two thirds of all brain regions met this criteria, underscoring the prevalence of rich club organization as a general principle of the brain network. By examining the extended rich club, we were able to gain a more detailed perspective on the potential role of the most peripheral regions in AD and MCI.

Gray matter atrophy was accelerated not only in the core rich club regions, but also in the extended rich club. This finding suggests that progressing gray matter atrophy in MCI and AD is generally associated with nodal degree and affects rich club properties at the network level. Progressive atrophy of rich club regions is consistent with previous work on network hubs in AD (Crossley et al., 2014; Dai et al., 2015) and confirms that cortical atrophy does not only co-localize with hub regions in general but also with the rich club in particular.

In healthy individuals, there was no significant difference in amyloid deposition between peripheral and extended rich club areas. However, we observed that the core rich club regions seemed to be less affected by amyloid deposition then all other regions. MCI and AD patients had a higher amyloid load than healthy controls, and the increasing amyloid deposition appeared to target rich club regions more extensively, confirming previous work (Buckner et al., 2009). The difference in amyloid deposition between core rich club regions and other brain regions observed in healthy individuals disappeared in both patient groups. Moreover, the extended rich club showed higher amyloid load than the periphery in MCI and AD patients. While we need to be cautious with our interpretation due to the lack of longitudinal data, these results support previous findings suggesting that disease progression might preferentially affect rich club regions (Yan et al., 2018).

On the functional level, we found more evidence for altered activation in peripheral regions than core rich club regions in AD and MCI across all meta-analyses. Even though the overall effect sizes were rather small, we can conclude that if functional differences occur in AD and MCI, they are more likely to target the periphery than the network’s structural core.

The identification of rich club atrophy, which has been observed in normal aging but appears to be accelerated in MCI and AD, raises questions regarding the underlying mechanisms and their relationship to amyloid pathology. Traditionally, atrophy has been considered a consequence of amyloid accumulation, as proposed by the amyloid cascade model, which posits an imbalance between amyloid-beta production and clearance as the initial event in the pathogenesis of AD (Jack et al., 2013, 2013). This imbalance leads to increasing amyloid deposits, triggering an inflammatory response that ultimately results in progressive atrophy and clinical dementia.

Interestingly, our study reveals that in healthy individuals, amyloid predominantly accumulates in the periphery of the brain, with a relative under-representation in the rich club regions. However, this distinction diminishes in individuals with MCI and AD, suggesting a shift in amyloid deposition towards the rich club. This observation supports the possibility that pathological proteins may spread trans-neuronally(Goedert, 2015; Prusiner, 1984), making the highly connected rich club regions susceptible to pathological processes originating in the periphery. Nonetheless, the question arises as to why significant atrophy occurs specifically in the rich club regions if amyloid deposition initially occurs in the periphery. Several scenarios can be considered to explain this phenomenon. Firstly, hub regions possess more gray matter volume compared to peripheral regions, rendering them more prone to faster volume loss in response to pathology. Additionally, hub regions have higher metabolic rates (Collin, Sporns, et al., 2014; Liang et al., 2013; Vaishnavi et al., 2010), which may amplify the detrimental effects of pathological processes. Furthermore, afferent projections from the periphery tend to converge onto hub regions (Harriger et al., 2012; Senden et al., 2018; van den Heuvel et al., 2012). Peripheral pathology could thus lead to diminished input and pronounce atrophy even beyond any local pathology resulting from increasing amyloid levels in the rich club (Fornito et al., 2015).

Although the precise computational role of the rich club remains a topic of ongoing investigation, emerging evidence suggests its potential involvement in synchronizing activity and influencing functional reconfigurations within the periphery (Senden et al., 2014, 2018). The observed differences in peripheral activation may therefore result from dysfunction within the rich club. Another plausible explanation for the observed peripheral activation differences lies in the activation of compensatory mechanisms. It has been suggested that plastic changes are more likely to occur in the periphery as the brain network adapts and reconfigures in response to AD-related pathology (Chen et al., 2020; Ma et al., 2022). Consequently, the functional deficits observed in the periphery could potentially represent compensatory mechanisms or early-stage adaptations, while functional impairments within the rich club regions may manifest at later stages of the disease. Quite typically, fMRI assessments involve early-stage patients who are still relatively less affected by the disease (Fox & Greicius, 2010). This supports the suggestion that the observed peripheral activation differences might indeed be indicative of compensatory processes at play in early-stage patients.

The suggested link between amyloid PET and structural atrophy in relation to the rich club is particularly noteworthy as it connects two major biomarkers for AD with the rich club (Jack & Holtzman, 2013). This is significant because previous studies on structural and functional connectivity have yielded inconsistent findings regarding potential alterations in the rich club during the pathogenesis of AD. Establishing a connection between the rich club and key biomarkers of AD is thus a major step towards understanding AD from the perspective of a disconnection syndrome.

This being said, the present results need to be interpreted with caution due to several notable limitations.

First, the cross-sectional nature of our study, involving PET and functional data, restricts the ability to draw conclusions about causal relationships and dynamic changes over time. Furthermore, the limited longitudinal data with only two data points for atrophy and the absence of conversions from healthy to MCI or from healthy to AD limit the interpretation of our findings, particularly in relation to disease progression. It is important to consider these limitations when interpreting, for instance, the PET findings. The observed significant differences in controls that diminish in MCI and AD raise the question of whether the suggested trend persists when additional data points at later disease stages are included.

Another limitation stems from the inclusion of different samples for different imaging modalities, which precludes a direct comparison of the molecular, structural, and functional aspects within the same individuals. This introduces the challenge of narratively linking the different levels of analysis. Furthermore, the lack of matching between samples regarding disease stages constrains direct comparisons between the different imaging modalities at specific disease stages and may introduce confounding variables that influence the observed effects.

On the other hand, a strength of our study is that we utilized independent data from elderly individuals and healthy subjects as a common reference to define the rich club. This approach ensures that the reported effects specifically relate to typical rich club regions of the healthy brain. There is evidence that the rich club reorganizes during normal aging (Grayson et al., 2013; Riedel et al., 2022) but also during AD (Cao et al., 2020; Ma et al., 2022). Choosing a common and independent reference ensures that our results are not confounded by remodeling and adaptation to disease processes. Additionally, we examined the extended rich club, which includes all regions within the rich club regime, not just the top degree nodes. By considering the extended rich club, we accounted for the inclusion of “next in line” regions that would become part of the narrow sense rich club over time. Since progressing atrophy and amyloid deposition in AD and MCI also involved the extended rich club, there is reason to assume that the observed effects are not confounded by qualitative changes to connectome organization in dementia and its prodromal stages.

In conclusion, while our study provides insights into the role of the rich club in AD pathogenesis, it is important to acknowledge the limitations associated with the cross-sectional design, limited longitudinal data, lack of conversions, and inclusion of different samples for different imaging modalities. These limitations should be carefully considered when interpreting the findings. Future research that addresses these limitations could further advance our understanding of the rich club and its implications in neurodegenerative diseases.

## Acknowledgement

We thank Melanie Grauer, Anna Neuburg, and Lea Staab for help in data organization. Data collection and sharing for this project was funded by the Alzheimer’s Disease Neuroimaging Initiative (ADNI) (National Institutes of Health Grant U01 AG024904) and DOD ADNI (Department of Defense award number W81XWH-12-2-0012). ADNI is funded by the National Institute on Aging, the National Institute of Biomedical Imaging and Bioengineering, and through generous contributions from the following: AbbVie, Alzheimer’s Association; Alzheimer’s Drug Discovery Foundation; Araclon Biotech; BioClinica, Inc.; Biogen; Bristol-Myers Squibb Company; CereSpir, Inc.; Cogstate; Eisai Inc.; Elan Pharmaceuticals, Inc.; Eli Lilly and Company; EuroImmun; F. Hoffmann-La Roche Ltd and its affiliated company Genentech, Inc.; Fujirebio; GE Healthcare; IXICO Ltd.; Janssen Alzheimer Immunotherapy Research & Development, LLC.; Johnson & Johnson Pharmaceutical Research & Development LLC.; Lumosity; Lundbeck; Merck & Co., Inc.; Meso Scale Diagnostics, LLC.; NeuroRx Research; Neurotrack Technologies; Novartis Pharmaceuticals Corporation; Pfizer Inc.; Piramal Imaging; Servier; Takeda Pharmaceutical Company; and Transition Therapeutics. The Canadian Institutes of Health Research is providing funds to support ADNI clinical sites in Canada. Private sector contributions are facilitated by the Foundation for the National Institutes of Health (www.fnih.org). The grantee organization is the Northern California Institute for Research and Education, and the study is coordinated by the Alzheimer’s Therapeutic Research Institute at the University of Southern California. ADNI data are disseminated by the Laboratory for Neuro Imaging at the University of Southern California.

